# Processing of oxidatively damaged DNA dirty ends by APE1

**DOI:** 10.1101/2021.11.28.470279

**Authors:** Amy M. Whitaker, Wesley J. Stark, Bret D. Freudenthal

## Abstract

Reactive oxygen species attack the structure of DNA, thus altering its base-pairing properties. Consequently, oxidative stress-associated DNA lesions are a major source of the mutation load that gives rise to cancer and other diseases. Base excision repair (BER) is the pathway primarily tasked with repairing DNA base damage, with apurinic/apyrimidinic endonuclease (APE1) having both AP-endonuclease and 3’ to 5’ exonuclease (exo) DNA cleavage functions. The lesion 8-oxo-7,8-dihydroguanine (8-oxoG) can enter the genome as either a product of direct damage to the DNA, or through polymerase insertion at the 3’-end of a DNA strand during replication or repair. Importantly, 3’-8-oxoG impairs the ligation step of BER and therefore must be removed by the exo activity of a surrogate enzyme to prevent double stranded breaks and cell death. In the present study, we characterize the exo activity of APE1 on 3’-8-oxoG substrates. These structures demonstrate that APE1 uses a unified mechanism for its exo activities that differs from its more canonical AP-endonuclease activity. In addition, through complementation of the structural data with enzyme kinetics and binding studies employing both wild-type and rationally designed APE1 mutants, we were able to identify and characterize unique protein:DNA contacts that specifically mediate 8-oxoG removal by APE1.

## INTRODUCTION

During times of oxidative stress, excess cellular reactive oxygen species (ROS) react with and modify the structure of nucleic acids both within the DNA and the free nucleotide pool^1–3^. These modifications induce mutations in the genome and are a major source of genomic instability that promotes multiple human disease, such as cancer. The base excision repair (BER) pathway has evolved as the cell’s primary defence against base lesions and single stranded breaks (SSB) that arise during oxidative stress (Figure 1A)^4–8^. The BER pathway consists of five major steps: (1) recognition and excision of a damaged base, (2) strand cleavage at the abasic site, (3) replacement of the excised DNA nucleotide, (4) processing of DNA ends, and (5) sealing of the DNA nick. Within this pathway, APE1 processes DNA damage with two distinct nuclease activities: AP-endonuclease (AP-endo) and 3′ to 5′ exonuclease (exo). APE1 cleaves the DNA backbone 5’ of the abasic site with its AP-endo activity. In contrast, the exo activity of APE1 proofreads and processes aberrant 3’ DNA termini that can prevent faithful repair, see red arrows in Figure 1A.

**Figure 1:**
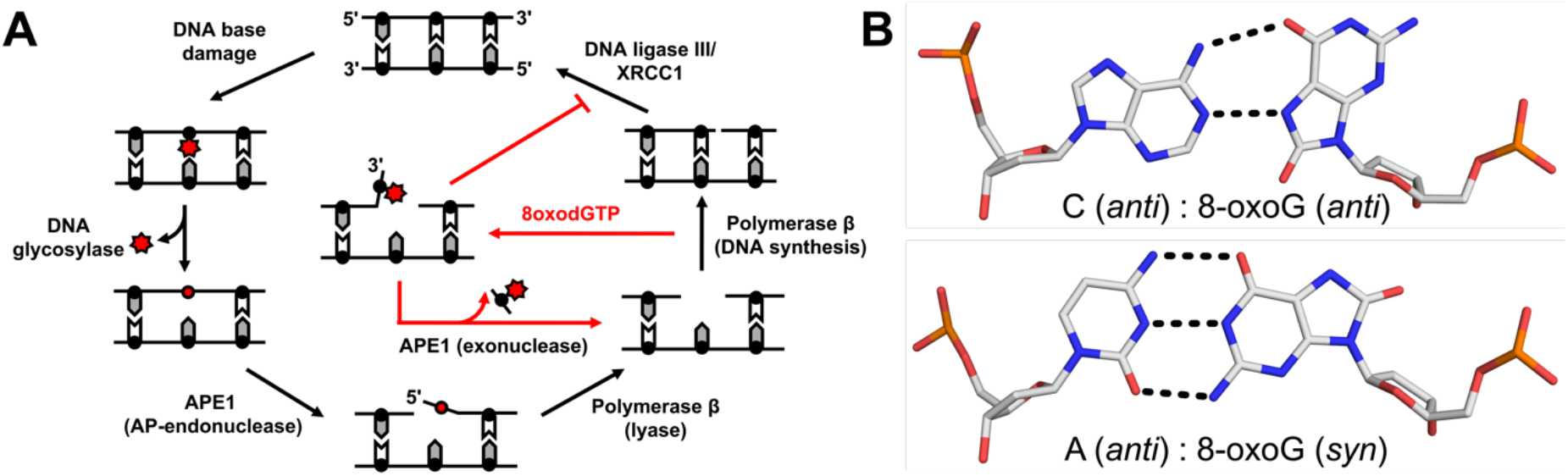
Base excision repair pathway. **A)** A schematic demonstrating how the BER pathway removes and replaces the oxidative lesion 8-oxoG. Red arrows highlight the role of the APE1 exo activity after 8-oxodGTP insertion by Pol !. **B)** The base pairings formed by 8-oxoG opposite A and C.

Oxidatively damaged DNA termini can be referred to as DNA “dirty ends” and require subsequent processing to repair the lesion and maintain genome stability^9^. One major oxidized nucleobase resulting from ROS is 8-oxo-7,8-dihydro-2’-deoxyguanosine, which is found in both DNA (8-oxoG) and in the nucleotide pool (8-oxodGTP)^1–3^. Even with cellular sanitizing activities, nucleotide pools contain enough 8-oxodGTP to promote mutagenesis^10,11^. Mutagenesis arises from the dual coding potential of the oxidative lesion, where 8-oxoG (*anti*) base pairs with cytosine (C) using its Watson-Crick face and 8-oxoG (*syn*) uses its Hoogsteen face to base pair with adenine (A), Figure 1B. Structural studies examining the insertion of 8-oxodGTP revealed that the modified nucleotide can escape general DNA polymerase discrimination checkpoints by modulating the highly charged DNA polymerase active site^3,12^. In addition, ROS can also attack the sugar moiety of DNA and directly produce nicks with damaged terminal ends^13^. These so-called ‘clustered lesions can result from ionizing radiation, which is deposited in small volumes of nanometer dimensions that produce ROS at high localized concentrations. Importantly, in either case, a DNA ligase would be responsible for sealing a nick with a dirty end. Unfortunately, 8-oxoG positioned at the 3’-end of a nick hastens abortive ligation and stabilizes the cytotoxic nick, thus resulting in persistent DNA containing SSBs with 3′-terminal 8-oxoG ends necessitating extrinsic end processing^3,14^.

APE1 has been shown to remove 3’-8-oxoG from nicked DNA substrates *in vitro*^13,14^. Moreover, cellular studies using whole cell extracts and immunoprecipitation experiments support a cellular role of APE1 in 3’-8-oxoG removal^13,15,16^. APE1 end processing of 8-oxoG, during which the oxidative DNA lesion is removed, results in a ‘clean’ 1-nt gapped substrate suitable for DNA synthesis by Pol β during BER (Figure 1A). Currently, the molecular mechanism used by APE1 for the end processing of the 8oxoG lesion is unknown. To probe this mechanism, we combined X-ray crystallography and enzyme kinetics to observe and characterize APE1 cleaning dirty DNA ends that arise from oxidative stress.

## MATERIAL AND METHODS

### DNA Sequences

The 21-mer nicked exonuclease substrate with a 3’-8oxoG used for crystallization was made using the following DNA sequences: opposing strand, 5′-GGATCCGTCGA(X)CGCATCAGC-3′, where X represents the base opposite O8; upstream strand with 3′ cytosine, 5′-GCTGATGCG(O8)-3′, where O8 represents the 8oxoGuanine lesion; downstream strand with a 5′-phosphate, 5′-TCGACGGATCC-3′. To generate the 30-mer nicked exonuclease substrates with a 3’-8oxoG for the kinetic and binding studies the following DNA sequences were used: opposing strand, 5′-GTGCGGATCCGTCGA(X)CGCATCAGCGAACG-3′, where X represents the base opposite O8; upstream strand with a 6-FAM (indicated by asterisk), 5′-*CGTTCGCTGATGCG(O8)-3′, where O8 represents the 8oxoGuanine lesion; downstream strand with a 5′-phosphate, 5′-TCGACGGATCCGCAT-3′. All sequences containing 8oxoG lesions were purchased from Midland Scientific, and all other sequences were purchased from IDT. Purified DNA substrates were annealed in buffer containing 50 mM tris and 50 mM KCl, and the concentration was determined by absorbance at 260 nm.

### Expression and Purification of APE1

Human wild-type and truncated APE1 (lacking 43 N-terminal amino acids) were expressed from pet28a codon optimized clones purchased from GeneScript. In order to generate the full-length and ΔAPE1 variants, site-directed mutagenesis was performed with a QuickChange II site-directed mutagenesis (Agilent). Once confirmation of correct sequencing was obtained, the plasmids were transformed into One Shot BL21(DE3)plysS E. *coli* cells (Invitrogen). Cells were growth in 2xYT at 37°C until OD_600_ was 0.6. The cells were then induced using isopropyl β-D-thiogalactopyranoside (IPTG) to a concentration to 0.4 mM. The temperature was then turned down to 20°C and cells were allowed to continue growing, while shaking overnight before being harvested. After harvesting, cells were lysed at 4 °C by sonication in 50 mM HEPES, pH 7.4, 50 mM NaCl, 1 mM EDTA, 1 mM DTT, and a protease inhibitor cocktail (1μg/mL leupeptin, 1 mM benzamidine, 1 mM AEBSF, 1 μg/mL pepstatin A). Lysate was clarified from cell debris by centrifugation for 1 hour at 24,424*g*. The pellet was discarded, and the resulting supernatant was passed over a HiTrap Heparin HP (GE Health Sciences) equilibrated with lysate buffer (50 mM HEPES, pH 7.4, 50 mM NaCl). APE1 was eluted from the column using a linear gradient of NaCl up to 600 mM. APE1 was then buffer exchanged into 50 mM NaCl and loaded onto a Resource S (GE Health Sciences) equilibrated in lysate buffer (50 mM HEPES, pH 7.4, 50 mM NaCl). APE1 was eluted from the column using a linear gradient of NaCl up to 400 mM. APE1 was subsequently loaded onto a HiPrep 16/60 Sephacryl S-200 HR (GE Health Sciences) equilibrated in 50 mM HEPES, pH 7.4, and 150 mM NaCl. SDS-PAGE was used to determine purity of the resulting fractions. Pure fractions were pooled together, and the final concentration was determined by NanoDrop One UV–Vis Spectrophotometer (Thermo-Scientific) at an absorbance of 280 nm. Protein was aliquoted and stored at −80 °C in a buffer of 50 mM HEPES, pH 7.4 and 150 mM NaCl.

### APE1 Crystallization and Structural Determination

For the 3’ 8oxoG structures three oligonucleotides, one with a 3’ 8oxoG, were used to form a were annealed in a 1:1:1 ratio to a final concentration of 2 mM using a PCR were used to form a 21-mer duplex with a central nick one with a central THF mimicking an AP-site, were annealed in a 1:1 ratio to a final concentration of 2 mM using a PCR thermocycler by heating for 10 min at 90°C and cooling to 4°C (1°C/min). For the A similar protocol was used to generate the 3’ mismatched DNA for the exonuclease complex crystals, in which. In both cases, the annealed DNA was mixed with C138A-ΔAPE1 to achieve a final concentration of 0.56 mM DNA and 10 - 12 mg ml^−1^ C138A-ΔAPE1. The single amino acid C138A mutation and truncation of the N-terminal 43 amino acids aid in crystallization^17,18^. For the AP-DNA complexes, product was generated by allowing APE1 to cleave the DNA in solution for 30 minutes on the bench prior to setting up crystallization trays. Vapor diffusion was utilized in order to crystalize the APE1:DNA complexes. The reservoir solution for crystallization of AP-endonuclease structures was 9 - 11% PEG 20K, 100 mM sodium citrate pH 5, and 200 mM magnesium chloride. For the exonuclease structures, the reservoir solution for crystal formation was 7 – 14% PEG 20 K, 100 mM sodium citrate, pH 5.0, 15% glycerol, and 5 mM calcium chloride. In all cases, crystals formed within a week at 20°C and were transferred to a cryosolution containing the mother liquor and 25% ethylene glycol. Data were collected at 100 K on a Rigaku MicroMax-007 HF rotating anode diffractometer equipped with a Dectris Pilatus3R 200K-A detector system at a wavelength of 1.54 Å. This allowed for anomalous data detection after phasing by molecular replacement with high redundancy. Data were processed and scaled with the HKL3000R software package^19^. Initial models were determined by molecular replacement with a modified version of a previously determined APE1–DNA complex (PDB 5DFF or 5WN4) as a reference. Refinement was carried out with PHENIX and model building with Coot^20,21^. The figures were prepared with PyMOL (Schrödinger LLC), and for simplicity only one conformation is shown for residues with alternate conformations unless otherwise noted.

### APE1 Exonuclease Activity Assays

APE1 exonuclease DNA cleavage activity was performed using a benchtop heat block set to 37 °C. DNA substrates used to measure APE1 incision activity contained a 5’,6-carboxyfluorescein (6-FAM) label and were designed with a centrally located nick flanked by a 3’-8oxoG and a 5’-phosphate. The reaction buffer was 50 mM HEPES, pH 7.4, 100 mM KCl, 3 mM MgCl_2_, and 0.1 mg ml^−1^ BSA. Final concentrations after mixing were 100 nM DNA substrate and 0 – 250 nM APE1, as indicated on the plots. After 30 minutes reactions were quenched by mixing with an equal volume of DNA gel loading buffer (100 mM EDTA, 80% deionized formamide, 0.25 mg ml^−1^ bromophenol blue and 0.25 mg ml^−1^ xylene cyanol). After incubation at 95 °C for 5 min, the reaction products were separated by 22% denaturing polyacrylamide gel. The indicated % product formation for each concentration of APE1 are the mean of three independent experiments. A GE Typhoon FLA 9500 imager in fluorescence mode was used for gel scanning and imaging, and the data were analyzed with Image J software^22^.

### Anisotropy Binding Studies

Fluorescence anisotropy measurements were used to quantify binding of wild-type APE1 to the FAM-labeled DNA substrates described above. Fluorescence anisotropy measurements were carried out on a Horiba FluoroLog Fluorimeter at 37 °C in a buffer consisting of 50 mM HEPES pH 7.4, 100 mM KCl, 5% (v/v) Glycerol, and 20 mM EDTA. The excitation and emission wavelengths were 485 and 520 nm, respectively, each with a 5 nm slit width. For all titrations, the concentration of the FAM-labeled DNA was 10 nM to stay below the KD and maintain sufficient signal to noise. Fluorescence anisotropy changes were normalized to the number of binding sites and fit to either a one-site binding model or to a two-site binding model with Kaleidagraph 4.5 (Synergy). A minimum of ten independent measurements were averaged for each APE1 concentration of the titrations to determine K_d_ values.

## RESULTS

### APE1 Engaged with Exonuclease Substrates Containing 3’-8oxoG Lesions

The 8-oxoG base has been shown to adopt a variety of conformations both within protein active sites and in duplex DNA. This arises from the dual coding potential of the 8-oxoG base, which can form stable base pairs with both cytosine utilizing its Watson-Crick face and adenine using its Hoogsteen face (Figure 1B). Moreover, DNA Pol β inserts 8-oxodGTP opposite A, forming a mutagenic base paring, and C, forming a non-mutagenic base pairing, with only minor discrimination^3^. To see how the APE1 active site accommodates 3’-8-oxoG, we crystallized APE1 bound with nicked, double stranded DNA substrates containing 8-oxoG at the 3’-end and a phosphate at the 5’-end of the SSB. These substrates represent potentially unligatable^14^, BER intermediates in which the repair polymerase, Pol β, has misinserted an 8-oxodGTP across from either a templating A or C. To obtain crystals of the substrate complexes, we employed a double-mutant catalytically dead variant of APE1 (E96Q/D210N)^3,23^. The resulting structures, Table 1, revealed the APE1 active site positioned to remove the 8-oxoG lesion from the 3′ end of the DNA nick using its exo activity. Both the overall structures (RMSD 0.255 Å) and the organization of the APE1 active sites around the 3’- 8-oxoG are similar between the two structures (Figure 2). The 3’-8-oxoG structures also revealed several key protein:DNA contacts. These regions include DNA intercalating residues, a hydrophobic pocket adjacent the 3’-8-oxoG, stabilizing contacts with the 5’-phosphate, and the catalytic residues (Figures 2B and 2D). Specifically, R177 and M270 intercalate the DNA by wedging between the 8-oxoG base and the opposing A or C, serving as a physical block to any potential hydrogen bonding between 8oxoG and the opposing base. Additional contacts are formed between the 5’ phosphate and the side chains of N222 and W280 and two stable water molecules. Moreover, importantly, the DNA backbone is in position for catalysis adjacent residues composing the catalytic triad (E96, N68, and D210).

**Table 1:** Data collection and refinement statistics.

|  | APE1/O8:A | APE1/O8:C |
| --- | --- | --- |
| **Data collection** |  |  |
| Space group | P2<sub>1</sub> | P2<sub>1</sub> |
| Cell dimensions |  |  |
| *a*, *b*, *c* (Å) | 71.11,66.49,91.98 | 71.21,65.19,91.33 |
| $\alpha$, $\beta$, $\gamma$ (°) | 90,110.69,90 | 90,110.26,90 |
| Resolution (Å) | 25-2.0 | 25-2.25 |
| *R*<sub>meas</sub> (%) | 0.145 (0.674) | 0.133 (0.660) |
| *CC*<sub>1/2</sub> | 0.829 | 0.239 |
| *I*/σ<sub>*I*</sub> | 15.04 (1.71) | 10.05 (1.72) |
| Completeness (%) | 98.6 (96.9) | 99.6 (98.4) |
| Redundancy | 3.7 (2.5) | 4.1 (2.2) |
| **Refinement** |  |  |
| Resolution (Å) | 1.99 | 2.24 |
| No. reflections | 96631 | 65330 |
| *R*<sub>work</sub>/*R*<sub>free</sub> | 0.2260/0.2649 | 0.1997/0.2452 |
| No. atoms |  |  |
| Protein | 4431 | 4403 |
| DNA | 859 | 857 |
| Water | 263 | 243 |
| B-factors (Å<sup>2</sup>) |  |  |
| Protein | 41.72 | 39.00 |
| DNA | 67.90 | 64.40 |
| Water | 42.69 | 39.64 |
| R.m.s deviations |  |  |
| Bond length (Å) | 0.008 | 0.009 |
| Bond angles (°) | 1.011 | 1.000 |
| PDB ID | 7SUV | 7SVU |

\*Highest resolution shell is shown in parentheses

**Figure 2:**
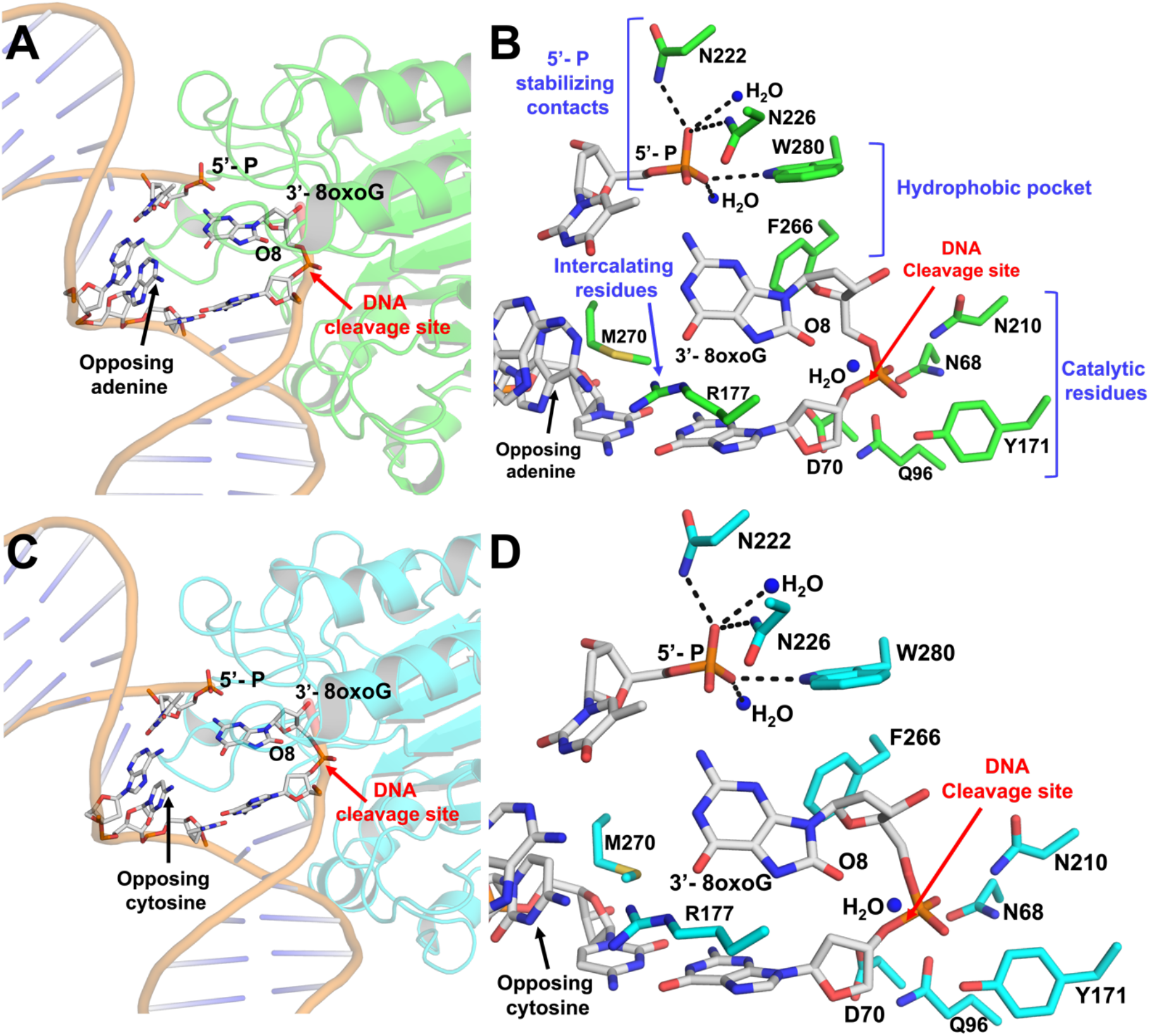
The APE1 active site engages 3’-8oxoG removal via its exo activity. **A)** Structural overview and **B)** active site configuration of APE1 (green) bound to a DNA exonuclease substrate containing a single-stranded nick with a 5’-P and a 3’-8oxoG opposing A. **C)** Structural overview and **D)** active site configuration of APE1 (cyan) bound to a DNA exonuclease substrate containing a nick with a 5’-P and a 3’-8oxoG opposing C. The DNA backbone is represented in orange cartoon and key DNA bases are represented as sticks in grey. Red arrows highlight the site of DNA cleavage.

Opposite either A or C, the 8-oxoG base adopts an *anti*-conformation, in which the Watson-Crick face of the base is oriented toward the major groove and the 8-oxoG base is accommodated via an open protein:DNA intra-helical binding pocket composed of hydrophobic residues F266 and W280, Figure 3. This contrasts with the endonuclease activity of APE1 with abasic DNA, of which the abasic site is base-flipped within the APE1 active site to adopt a catalytically competent conformation for cleavage during the AP-endo reaction^3,24^. The open active site accommodating 8-oxoG results from several structural variations between the 3’-8-oxoG exo and AP-endo structures, including a 10° sharper bend in the DNA and displacement of the backbone phosphate located 5’ of the nick. Of note, these structural differences are also observed in APE1 exo structures with a 3’-mismatched DNA substrate^4^.

**Figure 3:**
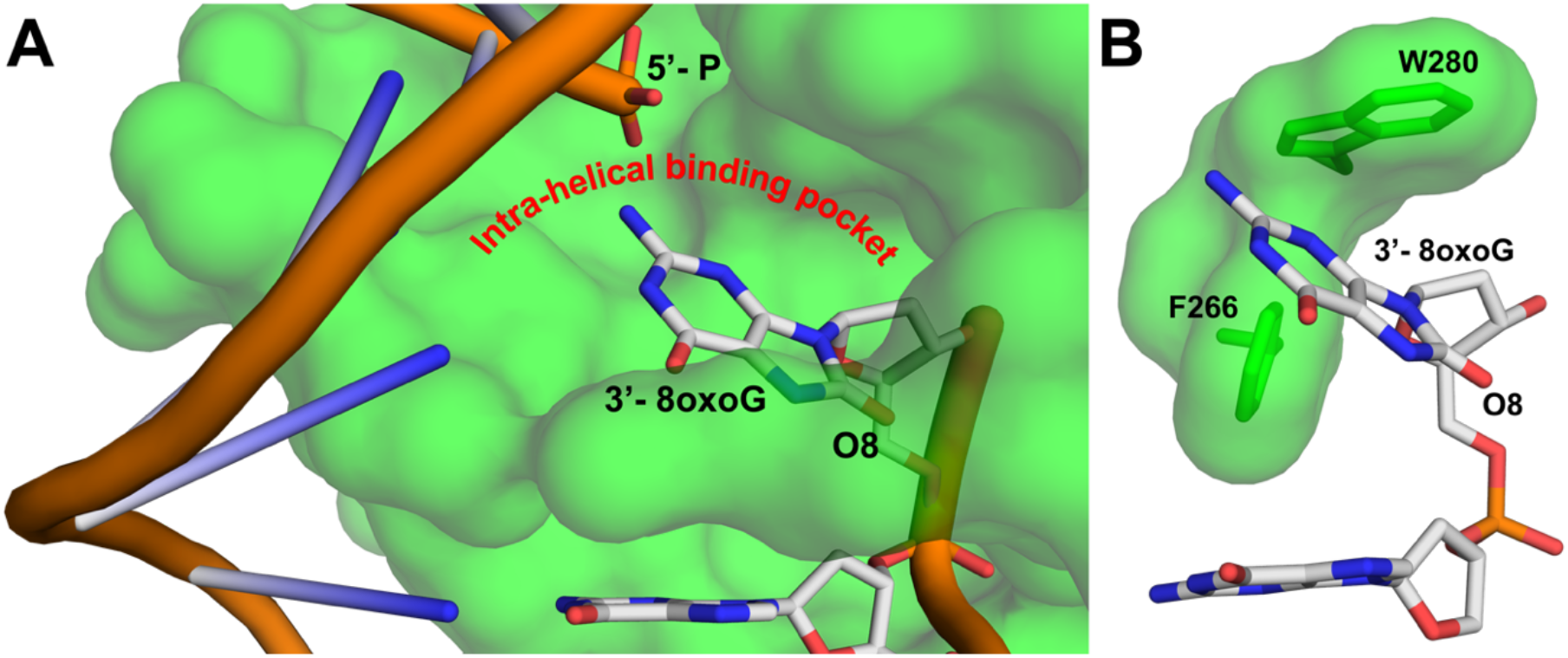
Key features of the APE1 exo reaction on 3’-8oxoG. **A)** The protein side chains R177 and M270 intercalate the DNA helix at the site of the nick, creating an intra-helical binding pocket containing the 3’-8-oxoG base. **B)** W280 and F266 compose the opposite side of the intra-helical binding pocket.

A closer look at the structure with 8-oxoG opposite A reveals the phosphate backbone oxygen (O5’) is coordinated to N174 (ND2) and N212 (ND2), Figure 4A. The 3’-OH of the sugar is also coordinated to N212 (ND2), while the two non-bridging oxygens are coordinated by N212 (OD1), N210 (OD1), Y171, and a water molecule. A second, ordered water molecule is in position to act as the nucleophile. This water is coordinated by the phosphate backbone oxygen (O5’) as well as one of the non-bridging oxygens, positioning the water atom 2.9 Å away from the phosphorus atom (Figure 4A). Moreover, the nitrogen atom of N210 (ND2) and the other active site water molecule are within H-bonding distance of the nucleophilic water at 2.7 and 3.1 Å, respectively. Also observed is a hydrogen bonding network between Q96, D70, N68, N210, and N212 that likely alters the pKa of N210 (D210 in WT APE1), thus facilitating the deprotonation of the water and subsequent nucleophilic attack at the phosphate backbone during cleavage. These contacts are consistent with a water mediated nucleophilic attack mechanism^24–26^. However, the structure lacks an anticipated bound active site metal ion, which is required for the APE1 exo activity^25,27^. This is most likely due to the APE1 D210N/E96Q active site mutations, which render a catalytically dead triad where Q96, N68, and N210 undergo moderate rotameric shifts to hydrogen-bond with one another^3^. This prevents E96Q from coordinating the metal, because NE2 is pointed toward the metal-binding pocket, and OE1 coordinates ND2 of N68. A similar phenomenon occurs with the N210 substitution, which coordinates OD1 of N68, thus resulting in ND2 coordinating the nucleophilic water molecule. Overall, the organization of the APE1 active site around the DNA substrate points to a unified mechanism for APE1 exo activity on 3’ mismatches and 3’ end damages including both the small oxidative lesion PG and the bulkier lesion 8-oxoG^4^.

**Figure 4:**
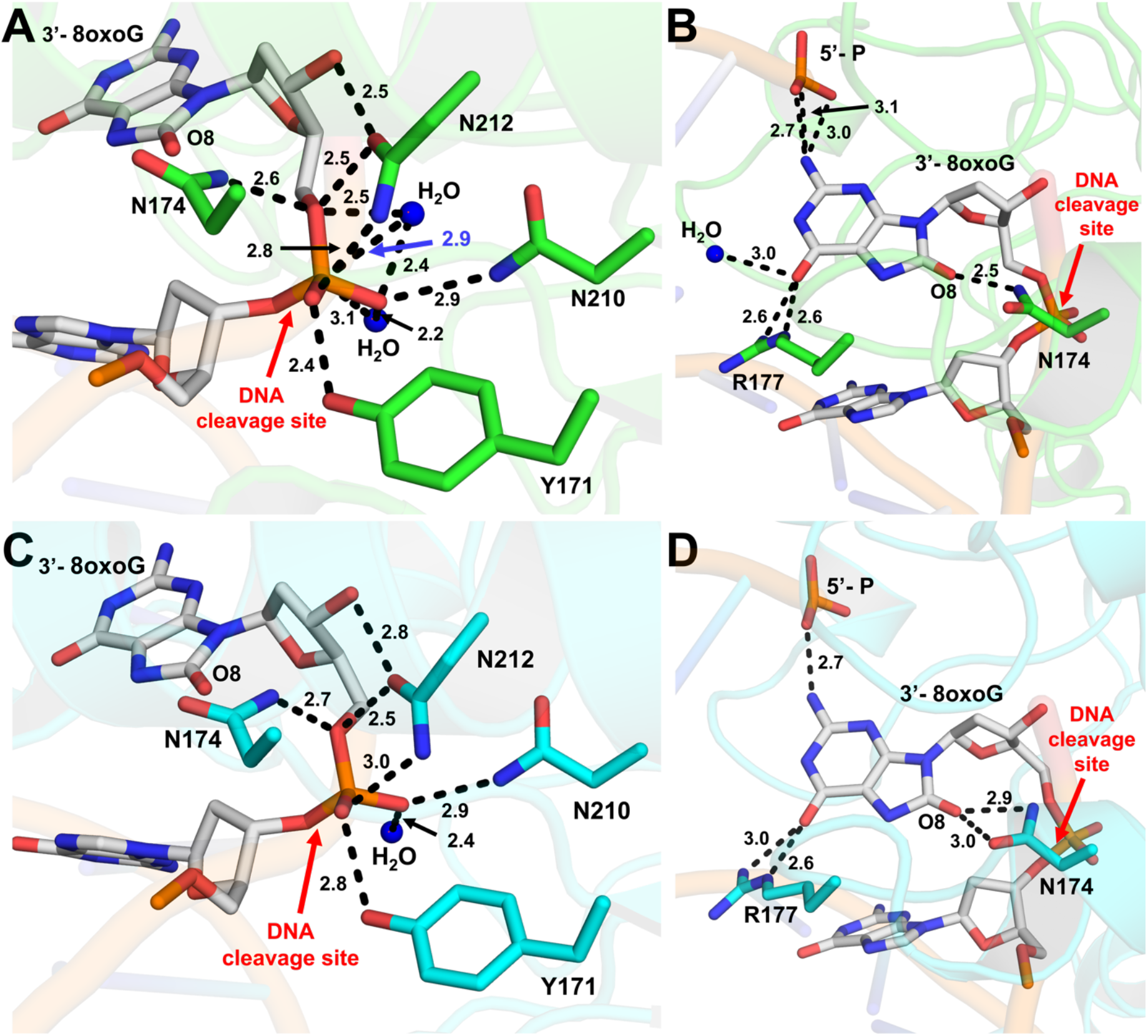
Interactions between APE1 and the 3’-8oxoG DNA substate. **A)** Contacts between APE1 protein side chains and the phosphate backbone of 3’-8-oxoG opposite A. **B)** Contacts between APE1 protein side chains and the 3’-8-oxoG base opposite A. **C)** Contacts between APE1 protein side chains and the phosphate backbone of 3’-8-oxoG opposite C. **D)** Contacts between APE1 protein side chains and the 3’-8-oxoG base opposite C. Key interactions are indicated by dashes and labelled with the distance in Å. O8:A is shown in green and O8:C is shown in cyan, the DNA backbone is represented in orange cartoon, and key DNA bases are represented as sticks in grey. Red arrows highlight the site of DNA cleavage.

In both structures, the 8-oxoG base is in position to hydrogen bond to the 5’-P and makes several contacts with the protein active site (Figures 4B and 4C). However, the precise contacts with the 8-oxoG base vary more substantially between the two structures than those between the backbone (Figure 4) likely due to its closer proximity to the variable opposing base, of which the double ring structure of A is larger than the single ring structure of C. Another similarity between the two structures is the intercalation of the DNA helix by R177, where it interacts with the Watson-Crick face oxygen (O6) of 8-oxoG using two of its side chain nitrogen atoms (NH2 and NE), Figure 4B and 4D). Of note, the O6 oxygen atom of 8-oxoG in the O8:A structure can additionally H-bond with a nearby stable water ion, Figure 4B. Positioned in between the two strands of the DNA helix, R177 is also in position to hydrogen bond with the opposing A or C base. These interactions between R177 and the opposing bases result in differences in its interactions with the 8-oxoG base. Specifically, the hydrogen bonding between R177 (NH2) and the 8-oxoG O6 oxygen atom is weaker, 3.0 versus 2.6 Å, in the case of the O8:C structure due to the contact between R177 and C requiring R177A to shift further towards the opposing strand than when it is opposite A (Figures 4B and 4D). Stabilizing the other side of the 8oxoG base, N174 is in position to form H-bonds with the lesion’s unique adducted oxygen atom (O8). N174 sits closer to the O8 of O8:A with 2.5 Å between it and the nitrogen (ND2) of the N174A sidechain, however in the case of O8:C both the N174 OD1 and ND2 are within hydrogen bonding distance at 2.9 Å and 3.0 Å, respectively (Figures 4B and 4D). The 8-oxoG bases of both structures also form H-bonds with the 5’ end of the DNA nick, with those in O8:A structure possibly being more extensive. These varying stabilizing interactions between the two structures is also reflected in the structural B-factors, which vary most substantially at the 3’-8-oxoG and opposing base positions (Supplemental Table 1). These differences are likely due to the shift in position of R177 between the two structures, which also pulls the base itself toward the opposing strand allowing it to maintain contact with both the opposing and 8-oxoG bases.

### Biochemical Characterization of APE1 and Variants with 3’-8-oxoG Substrates

To gain further insight into the substrate specificity of APE1 on 3′-8-oxoG substates, we examined the ability of APE1 to both bind and perform its exonuclease activity (3’-8-oxoG removal) on substates containing either A or C as the base opposing the 3’-8-oxoG (Figure 5). Importantly, these substates represent mutagenic (O8:A) and non-mutagenic (O8:C) base pairings. To determine the equilibrium binding affinities between APE1 and the 3’-8-oxoG substates, we measured fluorescence anisotropy of the DNA at increasing concentrations of APE1 from 0 to ~4000 nM and fit the resulting binding curve to a least squares fit non-linear regression (Figure 5A). The data indicate that APE1 binds the two substates, O8:A and O8:C, with similar affinities (K_d_) of 165.7 ± 31.38 nM and 205.6 ± 28.60 nM, respectively. To determine the relative exo activity of APE1 on these two substates, we next performed an APE1 activity assays. In these experiments, we observed APE1 product formation after 30 minutes over a range of APE1 concentrations ranging from 0.5 to 250 nM for each substrate. At each concentration of APE1, more product was observed for the O8:A substrate, opposed to O8:C, with about 2-fold more product formation for this mutagenic base pairing at the highest concentrations of APE1 (Figure 5B).

**Figure 5:**
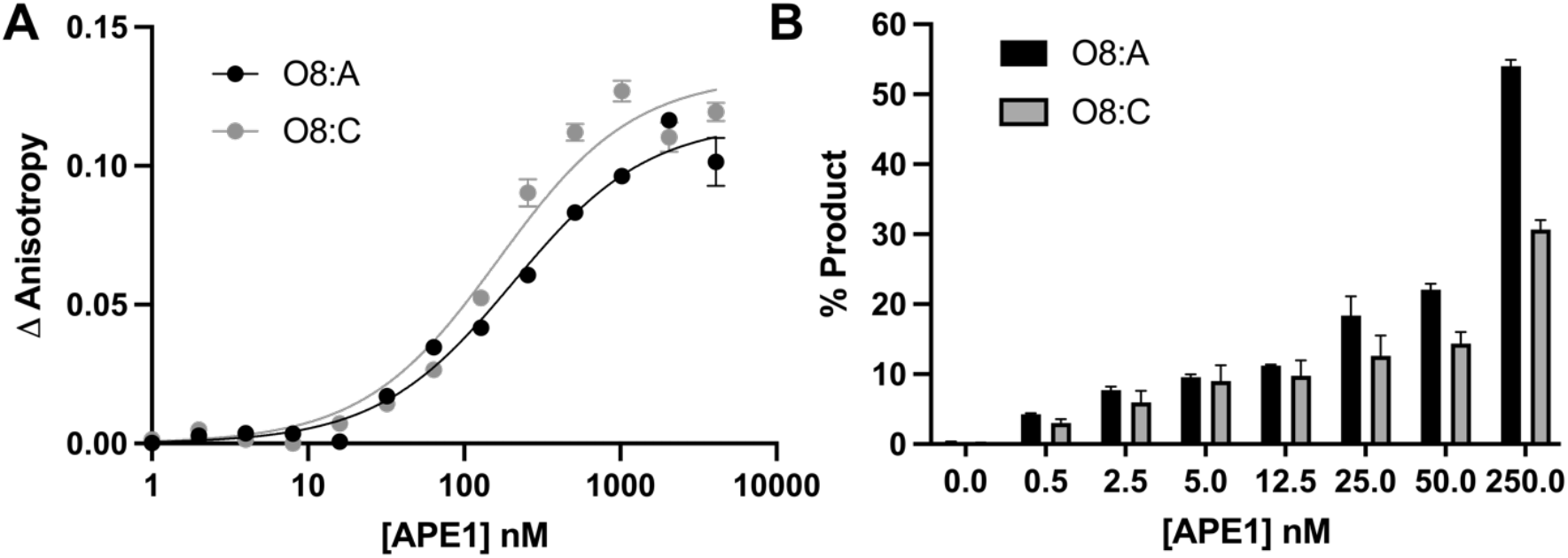
Functional characterization of the APE1 exo reaction on SSB 3’-8-oxoG, exo substrates. **A)** Binding of APE1 to the SSB 3’-8-oxoG substates with O8:A (black circles) and O8:C (grey circles). **B)** Formation of the APE1 5’ to 3’ exo cleavage product over a period of 30 min at 37°C for a range of WT APE1 concentrations from 0 to 250 nM APE1 with O8:A (black bars) and O8:C (grey bars) base parings at the 3’-end of the nick.

Previous work characterizing the exo activity of APE1 on a 3’-mismatched substrate demonstrated that mutating the residues that compose the enzymes hydrophobic pocket, W280 and F266, to alanine increases the steady-state rate of APE1 exo activity^24,28^ by 5-fold and 50-fold, respectively. A structure of the F266A APE1 variant engaged with a 3’-mismatched substrate indicated that the increase in activity was due to an expansion of the active site pocket, allowing the mismatched C to occupy an alternate, more catalytically active, conformation. The similarities between previous 3’-mismatched structures and the 3’-8-oxoG structures described above indicate that APE1 utilizes an analogous mechanism for its exo activity on these substrates as it does for 3’-mismatched substrates. To probe the role of these key active site pocket residues, we next utilized the APE1 cleavage activity assay on the same 3’-8-oxoG containing substates used above with W280A and F266A APE1 variant enzymes (Figures 6A and 6B). Our structures indicate that the side chains of both W280 and F266 constrict the size of the APE1 active site, and previous kinetic analyses indicate W280A and F266A APE1 mutants increase the exo catalytic rate^4,28^. In agreement, F266A showed a dramatic increase in the rate of product formation compared to WT APE1 for both O8:A (Figure 6A) and O8:C (Figure 6B). This indicates a universal role for F266 and the APE1 intrahelical binding pocket in APE1 exo activity. W280A, which modestly enhances exo activity in the case of a 3’-mismatch in comparison to WT APE1, on the other hand, modestly reduces the APE1 catalytic activity on exo substrates with a 3’-8-oxoG (Figures 6A and B). An overlay of the crystal structure of APE1 with 3’-mismatched C in the intrahelical binding pocket (PDB:5WN4) with those containing 3’-8-oxoG indicate a rotation in 8-oxoG of about 60° relative to the position of the 3’-mismatched C away from W280, perhaps explaining the different catalytic roles of W280 (Figure 6E). This rotation of the 8-oxoG base allows for the interaction between the adducted O8 oxygen and the side chain of N174.

**Figure 6:**
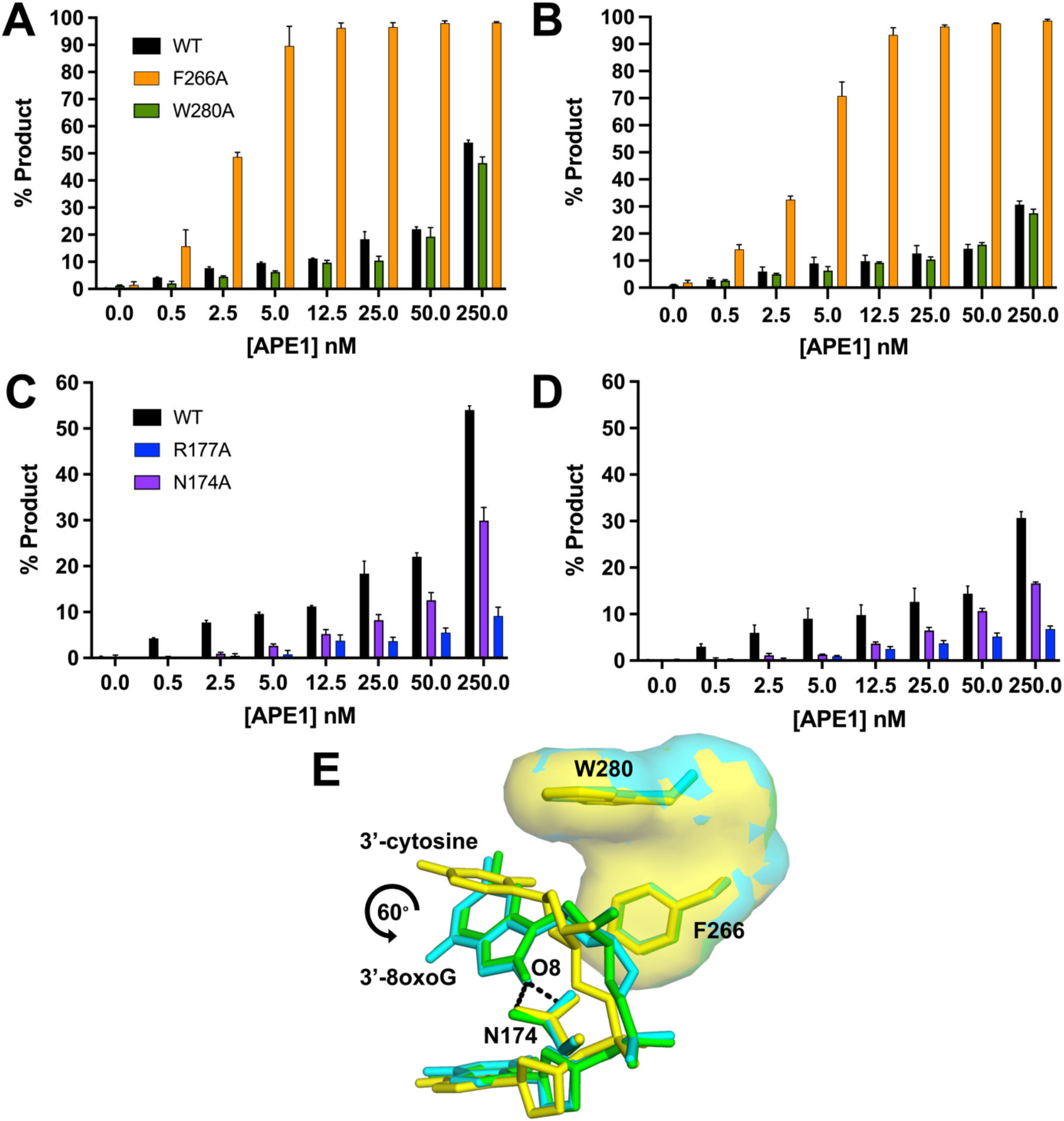
Functional characterization of the exo reaction of APE1 variants on nicked 3’-8-oxoG substrates. Formation of APE1 5’ to 3’ exonuclease product over a range of variant APE1 concentrations from 0 to 250 nM. **A)** compares WT (black), F266A (orange), and W280A (green) on a substrate containing 3’-8-oxoG opposite A. **B)** compares WT (black), F266A (orange), and W280A (green) on a substrate containing 3’-8-oxoG opposite C. **C)** compares WT (black), R177A (blue), and N174A (purple) on substrates containing 3’-8-oxoG opposite A. **D)** compares WT (black), R177A (blue), and N174A (purple) APE1 variants on a substrate containing 3’-8-oxoG opposite C. All assays were performed at 100 nM DNA, over a period of period of 30 min, at 37°C, and at the concentration of APE1 indicated on the legend. **E)** An overlay of the crystal structure of APE1 with a 3’-mismatched C (yellow) in the intrahelical binding pocket (PDB:5WN4) with those containing 3’-8-oxoG indicate a rotation in O8 of about 60° relative to the position of the 3’-mismatched C away from W280. O8:A is shown in green and O8:C is shown in cyan.

The 8-oxoG base makes unique interactions with the APE1 N174 and R177 side chains. To determine whether these interactions play a role in the APE1 exo activity on 3’-8-oxoG substates, we performed the same APE1 product formation assays with N174A and R177A APE1 variants (Figures 6C and 6D). Irrespective of the opposing base, N174 is in position to H-bond with the adducted oxygen of the 8-oxoG base (Figures 4B and 4D). When this interaction is lost, via mutation of N174 to an alanine, we observed APE1 to have reduced activity on both 3’-8-oxoG substates. R177 interacts with both opposing bases and the 8-oxoG base, and because the structure of the opposing bases (C vs A) varies, these interactions result in differences in the interactions between R177 and the 8-oxoG base. Moreover, R177 is thought to be an important residue for the product release step of APE1 cleavage, and when mutated to an alanine increases APE1 endo activity as well as its exo activity on a 3’-mismatch^3,4,29,30^. Strikingly, we observed R177A to have an opposite effect on 3’-8-oxoG substates, with dramatically decreased activity on both 3’-8-oxoG substates. This indicates that the R177 sidechain is critical for APE1 removal of the 8-oxoG lesion from the 3’-end of a DNA nick.

## DISCUSSION

Both oxidative DNA damage itself and its repair mediate the progression of many human diseases, including cancers, associated with genomic instability^17^. As a result, efficient BER plays a vital role in maintaining genomic integrity. On the other hand, BER failure is both cytotoxic and potentially mutagenic, thus contributing to the pathology of disease^17,31–33^ . Akin to genomic DNA, dNTPs are also vulnerable to oxidation by ROS, leading to the presence of their oxidized forms (e.g. 8-oxodGTP) in the nucleotide pool. During BER, oxidized nucleotide insertion by Pol β leads to ligation failure and the formation of a blocked 5’-adenylated, cytotoxic repair intermediate^14^. Importantly, because of the genomic instability resulting from their incorporation into the DNA during replication and repair, elevated levels of 8-oxodGTP and other oxidized nucleotides are associated with disease and cancer^34,35^. Moreover, because Pol β lacks a proofreading domain, misinsertions must be removed by a surrogate enzyme(s) to avoid genomic instability. Prior to this study, APE1 was shown to play this role, and can remove 3’-8-oxoG from nicked DNA substrates *in vitro*^13,14^. Moreover, cellular studies using whole cell extracts and immunoprecipitation experiments also support a cellular role of APE1 in 3’-8-oxoG removal^13,15,16^. However, the mechanism of APE1-mediated removal of 8-oxoG from the 3’-end of a DNA SSB remained elusive.

### The Conformation of 8-oxoG in the APE1 Active Site is Independent of the Opposing Base

One thing revealed by the substrate structures of APE1 engaged with a 3’-8oxoG, is that the 8-oxoG base adopts an *anti*-conformation independent of the nature (i.e. A or C) of the opposing base (Figures 2 and 3). This is distinct from the variable conformations of 8-oxoG observed when the base lesion is in duplex DNA, during which 8-oxoG(*anti*) base pairs with C and 8-oxoG(*syn*) adopts the Hoogsteen conformation to base pair with A^36^. The structural characterization of representative members of four families of DNA polymerases indicates that when 8-oxoG is in the flexible template binding pocket it can typically adopt either an *anti* or *syn* conformation depending on the nature of the incoming nucleotide as either C or A, respectively^37–47^. In contrast, as an incoming nucleotide, 8-oxodGTP usually favours the mutagenic *syn* conformation due to a lack of flexibility in the incoming nucleotide binding pocket and steric repulsion between O8 and its deoxyribose-phosphate^12,37,48,49^. Moreover, the insertion of 8-oxodGTP by Pol β has been characterized utilizing time-lapse X-ray crystallography^3^. This is particularly relevant, as it is this misinsertion of 8-oxodGTP by Pol β that proceeds its removal by the 3’ to 5’ exo activity of APE1. Interestingly, 8-oxoG insertion by Pol β across from C and A revealed 8-oxoG to be in either the *anti* or *syn* conformation during the reaction, respectively. However, with both templating bases, as Pol β reopened after catalysis the hydrogen bonding interactions between the bases were lost, and both adopted *anti*-conformations resembling those observed in the APE1 active site. This lack of stable base pairing between 3’-8-oxoG(*anti*) and either opposing base prior to APE1 engagement with the nicked DNA likely favours formation of the catalytically competent APE1:3’-8-oxoG(*anti*) conformations observed in our structures. In agreement with this, 3’-mismatches are removed by the APE1 exo activity more efficiently than matches^25,50^ and the efficiency of their removal is directly correlated to the thermostability of the mismatched base pair at the 3’-end of the nick^51^. Moreover, this also provides rationale for the apparent lack of strong discrimination depending on the opposing base, less than 2-fold (Figure 6).

### APE1 Utilizes an Intra-helical Hydrophobic Binding Pocket to Accommodate the 3’-8oxoG

While the above discussion addresses why there is not a large difference in APE1 activity depending on the nature of the opposing base, other factors must contribute to the modest difference in the APE1 exo activity observed between the two substates (Figure 5). To probe both these differences and furthermore to compare APE1 3’-8-oxoG removal with the other nuclease activities of APE1, we also characterized the 3’-8-oxoG exonuclease activities of a set of APE1 variants. Of the four variants, three (N174A, W280A, and R177A) resulted in reduced APE1 product formation while F266A, located in the intra-helical hydrophobic binding pocket, increased the rate of 3’-8-oxoG cleavage. The dramatic increase in APE1 exo activity on 3’-8-oxoG substrates in response to the F266A mutation (Figures 6A and 6B) was also observed previously in the case of 3’-mismatched substrates and was contributed to the mutation generating sufficient space in the APE1 active site for the 3’-base to adopt an alternative, more favourable, conformation^25,28^. Interestingly, F266A does not have the same effect on the APE1 AP-endo reaction and represents a separation of function mutant for APE1 exo and endo activities. The other hydrophobic pocket residue we tested, W280A, showed only a slight decrease in APE1 activity. Interesting, the opposite effect was seen in the case of 3’-mismatched exo substates^25^; however, in both cases the magnitude of change in response to the mutation was small, and it appears that the role played by W280 in APE1 exo reactions is minimal and is 3’-base-dependent in nature. Combined, the present study and previous work examining APE1 exo activity on 3’-mismatches indicate that the restrictive APE1 active site, and steric burden of large exo substrates (relative to an abasic site), are unifying features of the APE1 exo mechanism, and are likely responsible for the overall reduced incision rates for APE1 exo reactions compared to AP-endo reactions. Of note, the hydrophobic binding pocket comprised of F266 and W280 (Figure 3B), is not well conserved among APE1 homologs, and while it doesn’t appear to play a role in substate specificity between the various APE1 exo substrates, it does contribute to differences in substrate specificity between the 3’-exo and AP-endo activities of APE1 and the other Exo III family members^25,28^.

### Unique Contacts Between APE1 and the 8-oxoG Base Aid in its Removal

The N174A and R177A APE1 variants were selected to probe the functional relevance of the interactions between their native residue side chains and the 8-oxoG base. R177 is in position to H-bond with the opposing bases and the 3’-8-oxoG (Figures 2D,3B). On the other hand, N174 interacts with the unique O8 of the 8oxoG lesion. Both mutants decreased the APE1 cleavage activity on 3’-8-oxoG substates (Figures 6C and 6D), with the R177A mutation having a much more significant effect. This result was striking, as R177A has been characterized to have an opposite effect (albeit more modest), enhancing APE1 nuclease activity, with both AP-endo and 3’-mismatched exo substrates. This enhanced AP-endo activity of R177A APE1 has led to a model in which APE1 activity has been optimized for pathway efficiency by remaining bound to its incised product to facilitate ‘hand-off’ to the next enzyme in the pathway, Pol β^25,52^. Therefore, the role played by R177 appears to be unique in the case of 3’-8-oxoG, even among over exonuclease substrates.

Recent cell-based assays also explored the roll of the APE1 R177 sidechain using a complementation assay with APE1-deficient TM-Cre APE1^fl/fl^ MEFs and various APE1 mutants^53^. Interestingly, R177A APE1 was unable to rescue cell death. Moreover, the R177A mutant played a lesser role in protecting against MMS-induced cell death (which primarily generates abasic sites and alkylated damage) in comparison to its complementation efficiency under normal growth conditions. This may indicate that the role played by APE1 in removing the oxidative lesion 8-oxoG is particularly important for cell viability, however more work is needed to confirm this. Moreover, R177A was the only APE1 variant, including WT APE1, to have relatively the same activity on both 3’-8oxoG substrates, indicating that R177 and its interactions with the opposing base (A or C) may also be responsible for the differences in activity seen between the two substates. Therefore, R177A potentially serves as a separation of function mutation between the APE1 3’-8oxoG exo and AP-endo activities but also more specifically between its exo activities on 3’-8oxoG and its other exo activities, including 3’-mismatches.

### APE1 uses a Unified Mechanism for its Exonuclease Activities

APE1 is considered the major enzyme responsible for the removal of oxidatively damaged 3’-8-oxoG ends^54–58^ as well as 3’-end DNA mismatches from SSBs^50,59,60^. Here, by providing novel structural and kinetic data characterizing the APE1 3’ to 5’ exo activity on the 3’-8-oxoG lesions, we propose that APE1 uses a unified mechanism for its exo activities. The defining characteristics of this mechanism include DNA bending and fraying of the 3’ end at the site of the nick, intercalation of the DNA by protein sidechains R177 and M270 and positioning of the 3’-base within the intra-helical hydrophobic pocket. APE1 DNA bending is part of the larger DNA sculpting mechanism used by APE1 to engage its substates^61^. Supporting a unified mechanism for APE1 exo activity, APE1 similarly bends the DNA for both mismatched and 8-oxoG exo substates, but differently when bound to an abasic site (Figure 7A). The DNA intercalating residues R177A and M270 have been implicated in multiple roles during the APE1 nuclease activity, including facilitating protein:DNA interactions^24,25,29^, slowing product release (mentioned above)^3,4,24,29,30^, and more recently providing steric hindrance in the active site to distinguish exo substates^62^. Here we demonstrate that R177 plays different roles in the exonucleolytic cleavage of 3’-mismatches and 8-oxoG despite adopting similar conformations in the active sites of otherwise identical DNA substrates, Figures 7B through 7E. Instead, R177A seems to have unique 3’-base and opposing base interactions that result in the differences in activity observed between various exo substates.

**Figure 7:**
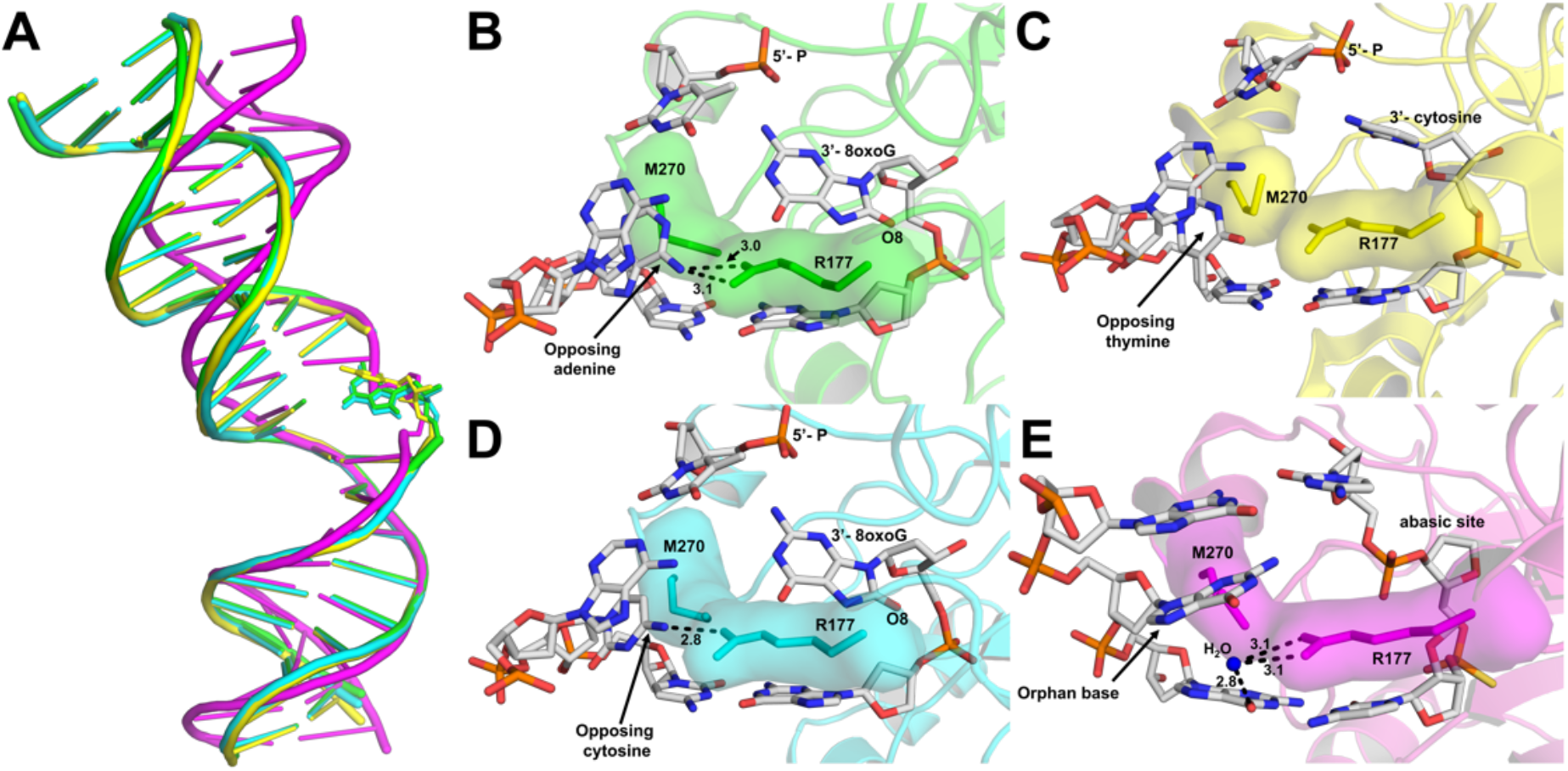
APE1 uses a unified mechanism for its exo activity. **A)** An alignment of the DNA from APE1:DNA complex structures highlighting the DNA bending, with 3’-8-oxoG:A in green, 3’-8-oxoG:C in cyan, 3’-mismatch in yellow, and abasic site DNA in magenta. The active sites of APE1 with **B)** 3’-8-oxoG:A, **C)** 3’-8-oxoG:C, 3’-mismatch, and **D)** abasic DNA. Key interactions are indicated by dashed lines and are labelled with the distance in Å, and key DNA bases are represented as sticks in grey.

Highlighting the importance of APE1 during DNA repair, polymorphisms and defects in APE1 have been identified in several human populations and are associated with the development of cancer^63–65^. To this point, APE1 represents a promising cancer therapeutic target^66,67^, making it essential to have a full understanding of the mechanisms of this multifunctional DNA repair enzyme. Moreover, the identification and characterization of separation of function mutants, such as F266A and R177A, is a key step in the rational design of molecular therapeutics that can specifically target select APE1 activities, including the 3’-8oxoG exo activity.

## DATA AVAILABILITY

Atomic coordinates and structure factors for the reported crystal structures have been deposited with the Protein Data bank under accession numbers 7SUV (O8:A) and 7SVB(O8:C).

## SUPPLEMENTARY DATA

**Supplemental Table 1:** B-factors for the 3’-8-oxoG exonuclease structures broken down by structural region. Red indicates those the most different between the two structures, yellow the second most, and green the least.

| B-factors | O8A (Å<sup>2</sup>) | O8C (Å<sup>2</sup>) |
| --- | --- | --- |
| all protein | 0.90 | 0.91 |
| R177 | 1.34 | 1.40 |
| W280 | 0.77 | 0.73 |
| N174 | 0.87 | 0.84 |
| F266 | 0.77 | 0.81 |
| all DNA | 1.46 | 1.50 |
| Opposing base | 1.31 | 2.00 |
| 5-Base (exo) | 1.16 | 1.34 |
| 3'-Base (exo) | 1.08 | 1.48 |
| overall | 1.00 | 1.00 |
| all water | 0.90 | 0.92 |

